# Toward Standardized Fucoidan Extractions from Brown Algae: Optimization of Under-Addressed Parameters to Maximize Yield and Purity

**DOI:** 10.64898/2026.11.17.767371

**Authors:** Jessica Carstens-Kass, Lauren K. Dedow, Siobhan A. Braybrook

## Abstract

Brown algal cell walls contain unique polysaccharides called fucoidans, which exhibit complex structural diversity and have applications across a range of industries. For these applications, fucoidan extraction methods should maximize yield while minimizing co-extracted impurities. However, extraction protocols vary widely, and their effects on yield and purity are rarely investigated. Thus, standardized and optimized fucoidan extraction methods are needed. Here, we aimed to identify optimal parameters for selected steps of a fucoidan extraction protocol. Starting with a commonly used protocol, we experimentally investigated three steps critical to yield and purity: (1) alcohol-insoluble residue preparation, (2) alginate removal, and (3) ion removal by dialysis. Our results show that sequential washes with 70% and 100% ethanol removed proteins, lipids, polyphenols, and pigments during alcohol-insoluble residue preparation without sacrificing fucoidan yield. We also demonstrate that extraction in calcium chloride solution reduces alginate-associated epitopes, while extraction in water does not. Finally, we show that dialysis with a membrane having a high molecular weight cut-off (12.4 kDa) results in fucoidan loss. Together, these findings inform specific recommendations for these three steps and provide a foundation for more standardized fucoidan extraction methods. Greater standardization could improve comparability across studies while facilitating the production of higher-yielding, purer fucoidan extracts for future research and downstream applications.

## INTRODUCTION

Fucoidans are a rare class of polysaccharides found only in brown algae (Phaeophyceae), sea urchins, and sea cucumbers. Fucoidans from sea urchins and cucumbers are structurally simple linear chains of repeating sulfated L-fucose sugar motifs (Pereira et al., 1999). On the other hand, fucoidans from brown algae are structurally complex; they can contain branches, non-fucose sugar monomers, and alternating linkage motifs (Pereira et al., 1999).

Fucoidans are a topic of research for a wide variety of reasons, ranging from brown algal biology to climate change to medicine. The functions of fucoidans in brown algae are not fully understood (Holtkamp et al., 2009), but they have been hypothesized to play a role in desiccation tolerance (Mabeau & Kloareg, 1987) and the structural integrity of the cell wall (Mabeau et al., 1990). In addition to their inclusion in the algal cell wall, fucoidans are secreted by brown algae into the ocean in significant amounts and are able to remain intact in the deep-sea sediment, thus acting as agents of carbon sequestration (Buck-Wiese et al., 2023; Vidal-Melgosa et al., 2022). Beyond their roles in the environment, fucoidans have bioactive properties including a plethora of therapeutic properties for humans and livestock (Abbas et al., 2025; Ale et al., 2011; Corino et al., 2021; Jin et al., 2014; Leal et al., 2025; Sanjeewa et al., 2021; Yu et al., 2024) as well as biostimulation of crops (Kaniyassery et al., 2024). Fucoidans also have unique material properties due to their polyelectrolytic nature, including ion conductivity in proton-exchange membranes (Vilela et al., 2020) and the ability to form bioactive multilayers for coating surfaces and capsules (Ho et al., 2015).

Yet, the fucoidan structures of individual brown algae are not well characterized and can vary based on taxonomy, environmental factors, and extraction methodology (Ale et al., 2011; Ko et al., 2026). The structure-function relationship of fucoidans is still unclear, leading to many outstanding questions, including: How does the structure of fucoidans affect their biological and ecological functions? Are certain fucoidan structures more likely to avoid microbial degradation and sequester carbon? How does the structure of fucoidans affect their bioactivities and material properties? Thus, to study the roles of fucoidans in nature and industry, it is necessary to be able to extract pure and native structures from brown algal tissue.

### Summary of Existing Fucoidan Extraction Techniques

Fucoidans account for 3-15% of the dry mass of brown algae (de Reviers, 1989; Mabeau et al., 1990; Percival & Ross, 1950). The rest of the dry mass is made primarily of other cell wall polysaccharides: alginate (40-60% (Mabeau & Kloareg, 1987)), cellulose (1.5-20% (Fangel et al., 2021)), and laminarin (3-35% (Dobrinčić et al., 2021)). Fucoidans must be separated from these other polysaccharides and from intracellular components like proteins, phenolics, lipids, and pigments to achieve the pure, native, extracts necessary for further studies. Fucoidan extraction protocols are varied, but many follow this common series of steps: (1) harvesting the tissue, (2) grinding the tissue, (3) washing the tissue with various solvents to remove proteins, phenolics, lipids, and pigments, (4) extracting fucoidans into an aqueous solution, (5) centrifuging, (6) dialyzing the supernatant to remove ions, and (7) lyophilizing to isolate the fucoidans (Figure 1). Each step requires consideration and optimization to maximize the purity and yield of the desired fucoidan molecules.

**Figure 1.**
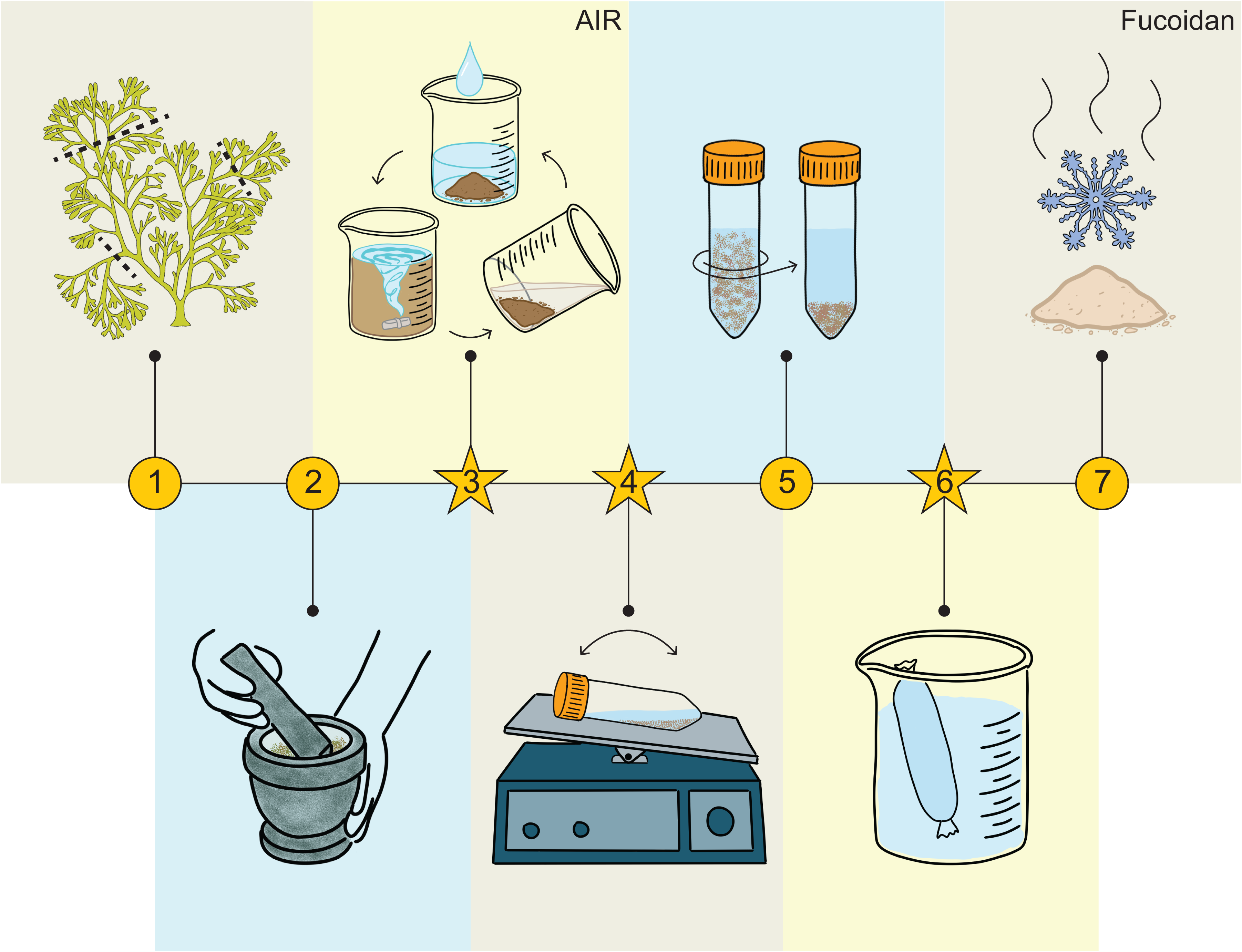
Schematic of the fucoidan extraction protocol. Stars indicate steps investigated in this study. 1) Harvest apical fronds of *Silvetia compressa* tissue. 2) Grind the tissue in liquid nitrogen to a fine powder. 3*) Wash the powder repeatedly with solvents to obtain the alcohol-insoluble residue (AIR). 4*) Incubate the AIR in CaCl_2_ solution (2% w/v) with mild agitation to cross-link alginates. 5) Centrifuge the mixture to separate the alginates (pellet) from the fucoidan solution (supernatant). 6*) Dialyze the fucoidan solution against deionized water to remove residual calcium and chloride ions. 7) Lyophilize the solution to obtain the final fucoidan product. Modified from an initial graphic created in BioRender (*BioRender*, n.d.) using Adobe Fresco (version 7.2.0).

#### Protein, polyphenolic, lipid, and pigment removal

Fucoidan extraction protocols historically have involved hazardous chemicals for the removal of impurities: lead acetate was used to precipitate proteins and alginate in 1950 (Percival & Ross, 1950), and live tissue was soaked in formaldehyde to remove phenolics in 1989 (de Reviers, 1989). Due to the risks associated with these chemicals, more modern protocols have focused on alternative chemicals for separating fucoidan from other cellular components.

Accordingly, more recent fucoidan extractions have removed impurities by washing with ethanol or acetone (Dobrinčić et al., 2020). These treatments produce what is commonly called an alcohol-insoluble residue (AIR), which is primarily composed of cell wall polysaccharides (Fangel et al., 2021). Only one study has measured the removal of impurities during AIR extraction, finding that two subsequent ethanol washes (both 80% or both 96%) achieved superior removal of polyphenols compared to the other wash schemes, while washing with acetone and then 96% ethanol achieved superior removal of pigments (Dobrinčić et al., 2021). The removal of lipids and proteins was not tested.

In addition to ethanol and acetone, other recent AIR extraction techniques have used methanol-chloroform solutions (Dobrinčić et al., 2020), which are more hazardous for the researcher and the environment. Methanol-chloroform-water solutions were originally used for lipid extraction from fish (Bligh & Dyer, n.d.), and were first applied in algae to extract bromophenolics from a red alga (Whyte & Southcott, 1970). Acetone (Dore et al., 2013) or acetone and methanol (Hentati et al., 2018) have been used in brown algal AIR extractions for the stated purpose of removing lipids and pigments; however, the lipid and pigment content before and after these wash steps has not been measured. In fact, the removal of lipids by a solvent has never been quantified in brown algae, to our knowledge.

In summary, it seems that ethanol, acetone, methanol, and chloroform are used in brown algal fucoidan extractions because the protocol for AIR extraction was carried over from plants, where specific combinations of these chemicals have been demonstrated to be the most effective (Fangel et al., 2021). However, impurity removal by each of these solvents has been only partially explored in brown algae, as explained above. In the present study, we aim to systematically determine which impurities (protein, polyphenols, lipids, pigments) are removed by which solvents (ethanol, acetone, chloroform-methanol solution) in AIR extraction from a brown alga.

#### Fucoidan extraction and alginate removal

After the removal of impurities, the residual AIR is stirred in an aqueous solution to extract the fucoidans. Mild solutions are preferred in order to avoid disrupting the native fucoidan structure. The most commonly used mild solutions for fucoidan extraction are water and calcium chloride solution (Dobrinčić et al., 2020). It has been suggested that CaCl_2_ solution may be better than water due to lower uronic acid (alginate) content in the resulting fucoidan product in two out of three species tested (January et al., 2019). This is expected, because calcium ions are known to cross-link with alginate, forming an insoluble precipitate (Beuder & Braybrook, 2023) that is separate from the water-soluble fucoidans. However, their conclusions are complicated by inconsistencies between the protocols at other steps. Here, we perform two fucoidan extractions using protocols that only differ in the solution used for the extraction step, and detect alginate in the fucoidan preparations to determine whether CaCl_2_ solution is superior to water for alginate removal during fucoidan extraction.

#### Ion removal

Finally, the fucoidan extract is often dialyzed to remove residual ions. These may be ocean salts that have made it through the AIR and fucoidan extraction steps, or calcium and chloride ions if CaCl_2_ solution was used for fucoidan extraction. Ion removal is particularly important if lyophilization is performed for fucoidan isolation in the subsequent step, because salts interfere with lyophilization. But, regardless of the isolation process, it is important to remove ions to obtain a pure fucoidan product for downstream utility. Dialysis membranes are available at a range of molecular weight cut-offs (MWCO), which specifies the molecular weight at which 90% of the macromolecular solute is retained by the membrane (Singh, 2015). MWCO specification is usually missing from fucoidan extraction methods, except in one study which specified the use of membranes with a 12-14 kDa MWCO (Nikolić Chenais et al., 2022). Fucoidan extracts contain fucoidans of a variety of sizes which are reported to range 25-5000 kDa (Neupane et al., 2020); however, these molecular weight analyses of fucoidans occur after a dialysis step, which possibly artificially increased the average molecular weight. In this study, we present the first investigation of how membrane MWCO affects fucoidan retention during dialysis.

#### Fucoidan isolation

Finally, the fucoidans must be isolated from the aqueous extraction solution. This is often accomplished via ethanol precipitation, because fucoidans are ethanol-insoluble (Dobrinčić et al., 2020). Alternatively, the fucoidan solution can be freeze dried if the researcher has access to a lyophilizer.

To better understand the impact of different fucoidan extraction protocol steps and their impact on the resulting purified fucoidan, we focused on 3 major areas: (a) AIR wash solvent combinations to maximize the removal of proteins, polyphenolics, lipids, and pigments, (b) CaCl_2_ incubation to demonstrate alginate removal, and (c) dialysis MWCO ranges to determine the correct range for maximizing the retention of fucoidan molecules. Figure 1 is a schematic representation of our fucoidan extraction protocol and the steps we investigated in this study.

## MATERIALS AND METHODS

### Sample collection and preparation

Specimens were collected in July of 2024 under California Department of Fish and Wildlife Scientific Collecting Permit GM-240580007-24058-001. *Silvetia compressa* (J. Agardh) E. Serrão, T. O. Cho, S. M. Boo et Brawley (Serrão et al., 1999) was collected from the southern end of the Fiscalini Ranch Preserve coastline in Cambria, CA, USA (rough coordinates: 35.548424, - 121.098468). Apical thallus tissue was harvested (Figure 1). The tissue was rinsed with freshwater and dried in an oven at 50°C for three days. The dried tissue was roughly blended in a food processor (Cuisinart Mini Prep Plus). The resulting granules were ground in liquid nitrogen with a basalt mortar and pestle to achieve a fine powder (Figure 1, Supplemental Figure 2).

### Cell wall extraction

To prepare the alcohol-insoluble residue, samples were washed with different combinations of the solvents ethanol, acetone, and chloroform:methanol (3:2 v/v). The following wash schemes were used: E70E100-five 70% washes, five 100% ethanol washes; E100-ten 100% ethanol washes; E70E100-A-five 70% washes, five 100% ethanol washes, two 100% acetone washes; E70E100-C-five 70% washes, five 100% ethanol washes, two 3:2 chloroform:methanol washes. 3 g of ground tissue was used for each wash scheme and all wash schemes were performed in triplicate. For each wash, the sample was stirred in a glass beaker with a magnetic stir bar, strong enough to create a vortex, with the solvent at room temperature for five minutes. The sample was then left for five minutes without stirring to allow the powder to settle. The solvent was decanted and the steps were repeated for the next wash (Figure 1). Decanted ethanol washes were retained for UV-visible light spectral absorbance quantification (see below). Samples were dried at 50°C overnight between the last 100% ethanol wash and subsequent acetone or chloroform-methanol washes, if applicable. The solvent-washed samples are expected to contain only cell wall polysaccharides and are referred to as the cell wall extract. The mass of each cell wall extract was measured (Supplemental Table 1).

### Fucoidan extraction

The entire mass of the cell wall extract was rocked in 20 ml of 2% CaCl2 solution (w/v) at room temperature for 30 hours. Samples were centrifuged at 39,800 x g for 10 minutes at 4°C. The supernatant was transferred to dialysis tubing (Spectra/Por® 3 Dialysis Membranes, MWCO 3.5 kDa, Avantor Sciences, Radnor, PA, USASpectra/Por 3RC). Replicate samples (n=3) were dialyzed against 2 L of ultrapure water (Milli-Q, Darmstadt, Germany) for 24 hours; water was replaced after 12 hours. Samples were transferred to 50-ml conical tubes and frozen at -80°C followed by flash freezing in liquid nitrogen. Immersing samples directly into liquid nitrogen was avoided as this resulted in splashing during freezing which interfered with formation of a ‘cake’ in the subsequent lyophilization step. Frozen samples were lyophilized using an Alpha 2-4 LSC Basic freeze-dryer (Martin Christ, Osterode, Germany) for 48-72 hours. In our hands, lyophilization results in a higher yield than ethanol precipitation (data not shown). The mass of each fucoidan extract was measured (Supplemental Table 1).

For the experiment comparing membranes with different MWCO, 10 ml of the extract was added to four types of dialysis tubing, in triplicate: 1 kDa (Spectra/Por® 7), 3.5 kDa (Spectra/Por® 3), 6-8 kDa (Spectra/Por® 1), and 12,400 kDa (Sigma-Aldrich, Product #D0655).

### L-fucose quantification

The L-fucose assay kit (MegaZyme, Bray, Ireland) was performed following the manufacturer’s instructions for polysaccharide and fibrous plant material, with modifications as follows: 2-5 mg of sample were added to a 2-ml polypropylene microcentrifuge tube with 1 ml 1.3 M hydrochloric acid. Samples were vortexed to mix and incubated in a heating block at 95°C for one hour, with burping to release pressure after the first 15 minutes. Samples were cooled to room temperature and neutralized with 1 ml 1.3 M sodium hydroxide. Water was added to bring the sample concentration to 1 mg/ml. Samples were centrifuged at 1500 x g for 10 minutes.

A standard curve of L-(-)-fucose (Sigma-Aldrich, St. Louis, MO, USA Product #F2252) was prepared at concentrations of 0-800 μg/ml, in increments of 200 μg/ml. In a 96-well plate, 10 μl of sample or standard solution was added along with 200 μl water, 40 μl buffer solution, and 10 μl NADP^+^ solution. The following shaking, incubation, and measurement steps were carried out on a SpectraMax iD3 microplate reader (Molecular Devices, San Jose, CA, USA). The plate was shaken orbitally at medium intensity for 30 seconds with the lid on, and then incubated at 37°C for 4 minutes with the lid off. The absorbance at 340 nm was measured. 5 μl of L-fucose dehydrogenase suspension was added. The shaking step was repeated and the plate was incubated at 37°C. The absorbance at 340 nm was measured after 10 min.

The L-fucose content of the standards was plotted against the ΔA340 to generate a linear regression equation. For experimental samples, this equation was used to convert ΔA340 to L-fucose content. These data and calculations can be found in Supplemental Tables 2, 8, and 9.

### Sulfate quantification

Sulfate was quantified by the Azure A method as described by Torode et al. (Torode et al., 2015) with modifications. Sample solutions were prepared at 200 μg/ml. In the absence of an ideal standard (sulfated carbohydrate with known sulfate content), a standard curve of commercial crude fucoidan from *Fucus vesiculosus* (Sigma-Aldrich, Product #F5631) was prepared at concentrations of 0-600 μg/ml. In a 1-ml cuvette, 50 μl of sample or standard solution was mixed with 50 μl 2.5 M sulfuric acid. 900 μl of 20 μg/ml Azure A dye solution was added to the cuvette and mixed. The absorbance was read at A620 using a Nanodrop One^C^ (ThermoFisher Scientific, Waltham, MA, USA), using water as a blank. Azure A dye binds to sulfate in solution, causing an immediate and proportional color change from blue to purple; decreased absorbance at 620 nm (blue) indicates sulfate in the sample.

To estimate the actual weight of sulfate in the sample, it was assumed that the standard and sample fucoidans were homofucans with a fucose:sulfate ratio of 1:1, based on the reported ratios in *Fucus* species: 1:1 in *Fucus serratus* (Bilan et al., 2006) and 1:1.21 in *Fucus distichus* (Bilan et al., 2004). A monosulfated fucose monomer is 33% sulfate by weight in atomic mass units (Supplemental Figure 3B, (0.086 ÷ 243.209)(100) = 33%). Thus, 42% of the mass of the standard used in the experiment was plotted against the A620 to generate a linear regression equation. For experimental samples, this equation was used to convert A620 to sulfate content. These data and calculations can be found in Supplemental Tables 3 and 10.

### Protein quantification

Protein was quantified using the Bio-Rad Protein Assay (Bio-Rad Laboratories, Hercules, CA, USA) with modifications. A standard curve of bovine serum albumin was prepared at concentrations between 100-900 μg/ml, in 100 μg increments. Experimental sample solutions were prepared at 15 mg/ml. To a cuvette, 20 μl of each standard or sample solution was added along with 1 ml Bradford Reagent. Samples were mixed by aspirating with a pipette and incubated at room temperature for 10 minutes. The absorbance at 595 nm was measured using a Nanodrop One^C^.

The protein contents of the standards were plotted against the A595 to generate a linear regression equation. For experimental samples, this equation was used to convert A595 to protein content. These data and calculations can be found in Supplemental Tables 4 and 11.

### Polyphenol quantification

The Folin-Ciocalteu assay was performed as described by Parys et al. (Parys et al., 2007) with modifications. A standard curve of phloroglucinol was prepared at concentrations of 0-20 μg. Sample solutions were prepared at 200 μg/ml. 800 μl of standard or sample solution and 50 μl of Folin and Ciocalteu’s phenol reagent 2 M (Sigma-Aldrich, Product #F9252) were added to a 1 mL cuvette. The reactions were mixed by aspirating with a pipette and incubated at room temperature for five minutes. 150 μl of 20% sodium carbonate solution (m/v) was added, producing an immediate color change upon reaction. Samples were mixed by aspirating with a pipette and incubated for 40 minutes in the dark. The absorbance at 760 nm was measured using a NanoDrop One^C^.

The polyphenol contents of the standards were plotted against the A760 to generate a linear regression equation. For experimental samples, this equation was used to convert A760 to polyphenol content. These data and calculations can be found in Supplemental Tables 5 and 12.

### Lipid quantification

A 5 mg/ml solution of cholesterol was prepared in 2:1 (v/v) chloroform-methanol solution, and a standard curve was prepared by adding 4-28 μg of cholesterol to tubes in increments of 4 μg. Sample solutions were prepared at 15 mg/ml in water. The remainder of the sulfo-phospho-vanillin assay protocol was carried out exactly as described by Zhong et al. (Zhong, 2016). The lipid contents of the standards were plotted against the ΔA540 to generate a linear regression equation. For experimental samples, this equation was used to convert ΔA540 to lipid content. These data and calculations can be found in Supplemental Tables 6 and 13.

### UV-visible light spectral absorbance quantification

Solutions of fucoidan samples were prepared at 15 mg/ml. The commercial fucoidans used were as follows: fucoidan from *Macrocystis pyrifera*, >85% pure (Sigma F8065); fucoidan from *Fucus vesiculosus*, crude (Sigma F5631); fucoidan from *Fucus vesiculosus*, >95% pure (F8190); and fucoidan (species and purity not specified; Thermo Scientific 467460010). Solutions of commercial fucoidans were also prepared at 15 mg/ml, except for F5631, which was prepared at 2 mg/ml due to limited materials. Normalization of the data is expected to correct for the difference in concentration. The UV-visible light spectral absorbance of the sample solutions and the ethanol supernatants (see above) was measured using a Nanodrop One^C^ (Supplemental Tables 14 and 15). For the ethanol wash supernatants, we multiplied the measured absorbance of 1 ml of supernatant by 1000, because 1 L of ethanol was used in total for the 10 wash steps. For the fucoidan samples, we divided the measured absorbance of a 15 mg/ml sample by 15 and multiplied the quotient by the total sample mass (82.4 mg for E100 and 71.1 mg for E70E100).

### Alginate detection

Sample solutions were prepared at 1 mg/ml in PBS. Alginate epitopes were detected through an enzyme-linked immunosorbent assay (ELISA) using alginate-directed antibodies as described by Torode et al. (Torode et al., 2016), with slight modifications. A NUNC Maxisorp 96-well plate (ThermoFisher Scientific) was coated with 100 μl/well of sample solution, or PBS as a blank, and incubated at 4°C for 4 hours with rocking. Wells were washed thrice with 200 μl ddH2O to remove unbound antigen. Wells were incubated with 200 μl blocking solution (5% milk in PBS) at 4°C overnight with rocking. Wells were washed thrice with 200 μl ddH2O, then incubated with 100 μl brown algal monoclonal antibody (BAM) 8 (1:25 dilution) or BAM10 (1:200 dilution) in 5% milk in PBS for 1 hour at 4°C, rocking. Wells were washed thrice with 200 μl ddH2O, then incubated with 100 μl anti-rat horseradish peroxidase-conjugated antibody (1:1000 dilution) in 5% milk in PBS for 1 hour at 4°C, rocking. Wells were washed thrice with 200 μl ddH2O. Wells were incubated with 100 μl of tetramethyl benzidine (TMB) substrate (0.1 M sodium acetate buffer, pH 6.0, 0.1% TMB, 0.009% (v/v) H2O2). The plate was incubated in the dark for 20 minutes to allow for color development. The reaction was stopped by the addition of 100 μl 1 M sulfuric acid solution. The absorbance was read at 450 nm. The raw data can be found in Supplemental Table 16.

### Carbohydrate gel electrophoresis

Fucoidan samples made with dialysis membranes of different MWCO (see above) were prepared as 2 mg/ml solutions. Three volumes of ethanol were added to each sample solution, and samples were centrifuged at 20,817 x g for 2 minutes to precipitate fucoidans. The supernatant was discarded and tubes were inverted on a paper towel to dry.

The carbohydrate polyacrylamide gel electrophoresis (C-PAGE) method was adapted from the methods described by Goubet et al. and Gonzalez et al. (Gonzalez et al., 2024; Goubet et al., 2011). Dried pellets were resuspended in 5X C-PAGE sample buffer (0.03% alcian blue, 25% glycerol, in 1M Tris-borate, pH 8.2). The alcian blue was included in the sample buffer to help with sample visualization while loading the gel. 500 μg of each sample were run through a 0.5% agarose gel (15 x 15 cm) at 100 V for 1 hour, with 1M Tris-borate, pH 8.2 as running buffer. The gel was stained with 0.1% toluidine blue dye in a 3% acetic acid, 50% ethanol (pH 2.5) solution for 1 hour, then destained with a solution of 20% methanol, 10% acetic acid. The gel was placed on a light box and imaged with the CamScanner app on an iPhone. Image contrast was enhanced in Apple Preview. The signal intensity of each lane was quantified using the Plot Profile tool in Fiji (Schindelin et al., 2012). The raw data can be found in Supplemental Table 17.

### Data visualization

Plots were generated with Matplotlib (Hunter, 2007).

## RESULTS

### Protein and polyphenolic removal is enhanced by a combination of 70% and 100% ethanol washes

Sequential solvent washes are common practice in cell wall studies in plants (Fangel et al., 2021) and were of particular interest to us for use in brown algae. Ethanol is the most common and safest solvent for AIR extraction (Figure 1 step 3), but it is used at varying concentrations (usually 80-95%) without explanation (Dobrinčić et al., 2020). For our study design, we took inspiration from Torode et al., who performed washes with sequentially increasing ethanol concentrations of 70, 80, 90, and 100% (Torode et al., 2015). Since fucoidan is water-soluble, we hypothesized that lower-percentage ethanol solutions may lead to fucoidan loss, but may also facilitate the penetration of the solvent into the cell wall material, allowing for better impurity removal. To test the effect of ethanol solution concentration in AIR extraction from brown algae, we prepared *Silvetia compressa* samples and treated them with two wash schemes: 1) 10 washes with 100% ethanol (E100); and 2) 5 washes with 70% ethanol followed by 5 washes with 100% ethanol (E70E100; Figure 2A). We then compared the yield and composition of each sample.

**Figure 2.**
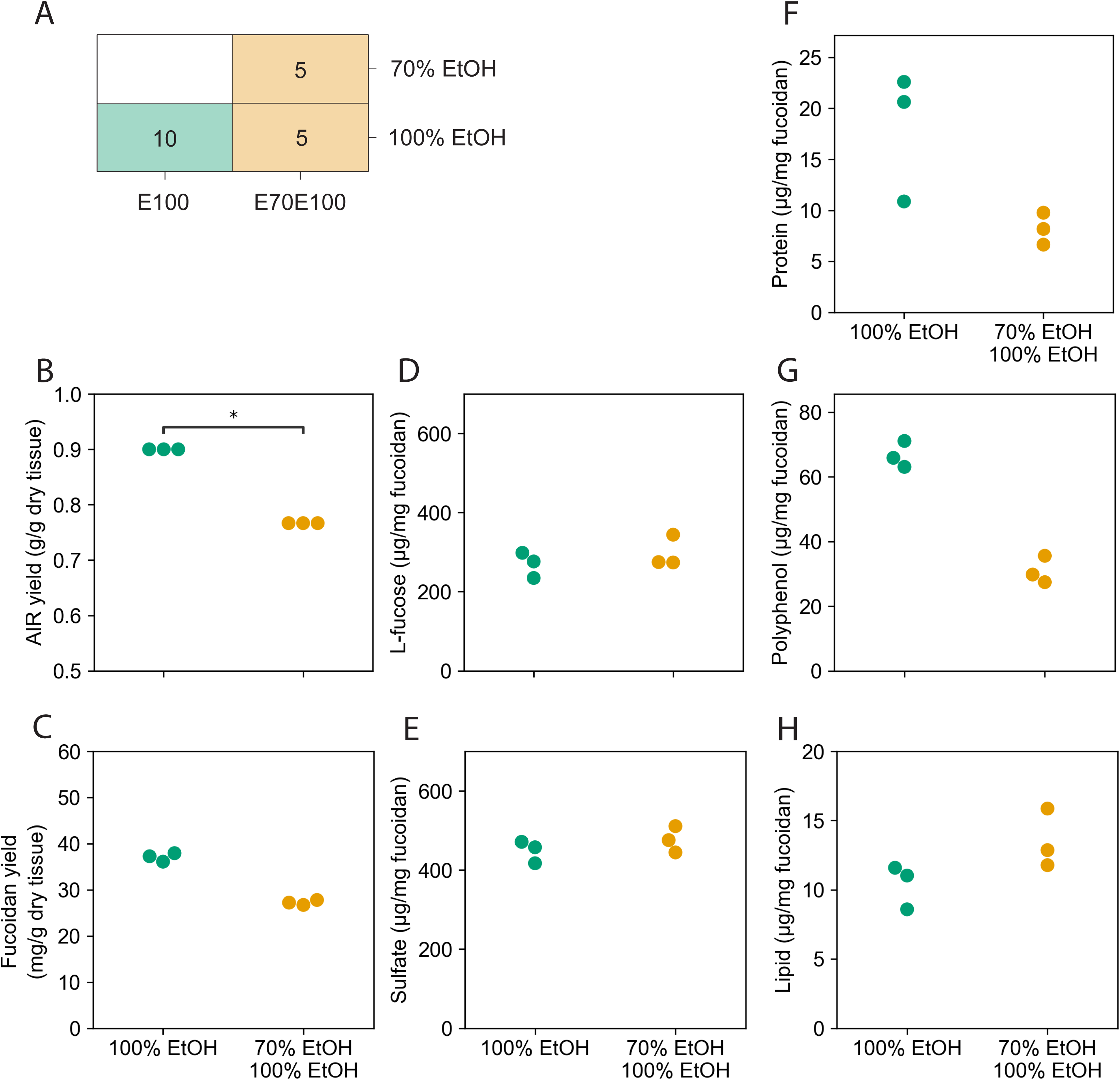
70% ethanol washes improve impurity removal from fucoidans. A) Schematic of the solvent wash schemes. The numbers inside the shaded boxes indicate the number of 5-minute washes that were performed with the solvent indicated on the right. B) AIR mass per gram of dry tissue for each wash scheme. C) Fucoidan mass per gram of dry tissue for each wash scheme. D) L-fucose content per mg of total fucoidan as measured by an enzymatic L-fucose assay kit (MegaZyme). E) Sulfate content per mg of fucoidan as measured by the Azure A assay. F) Protein contamination per mg of fucoidan as measured by the Bradford assay. G) Polyphenol contamination per mg of fucoidan as measured by the Folin-Ciocalteu assay. H) Lipid contamination per mg of fucoidan as measured by the sulfo-phospho-vanillin assay. Statistical significance was calculated using the Mann-Whitney-Wilcoxon test two-sided, * p ≤ 0.05.

L-fucose is the defining sugar monomer of fucoidan molecules, and the L-fucose sugars are often sulfated. To estimate the amount of actual fucoidan in the extracted fucoidan samples (Figure 1 step 7), we quantified the L-fucose content using an L-fucosidase enzyme-based assay kit, and the sulfate content using the Azure A assay (Torode et al., 2015). Then, to quantify the impurities in the extracted fucoidan samples, we quantified the protein content using the Bradford assay (Bradford, 1976), the polyphenol content using the Folin-Ciotalteu assay (Parys et al., 2007), and the lipid content using the sulfo-phospho-vanillin assay (Zhong, 2016).

The E100 wash scheme yielded significantly more AIR mass per gram of dry weight compared to the E70E100 wash scheme (Figure 2B). The higher AIR yield may indicate that more fucoidans were retained or that fewer impurities were removed. The E100 wash scheme yielded slightly more fucoidan mass per gram of dry weight (Figure 2C), but the fucoidan contained similar amounts of L-fucose and sulfate compared to the E70E100 samples (Figure 2D and E), indicating that the higher AIR yield was due to higher impurity content rather than higher fucoidan retention. Indeed, the fucoidans produced by the E100 wash scheme contained more protein and polyphenol impurities (Figure 2F and G), which likely account for part of the higher AIR and fucoidan mass. The fucoidans produced by the E100 wash scheme unexpectedly contained slightly less lipids than the fucoidans from the E70E100 wash scheme, but the lipid content was very low for both samples, suggesting that there is a low concentration of lipid impurities in the tissue to begin with. Taken together, these data indicate that the combination of 70% and 100% ethanol washes is superior for protein and polyphenol removal compared to washing with only 100% ethanol, leading to an overall purer product, without decreasing fucoidan yield.

### 70% and 100% ethanol washes also remove more pigment contamination

In addition to proteins, polyphenols, and lipids, ethanol is reported to remove pigments during AIR extraction (Figure 1 step 3). To visualize the removal of pigments, we measured the ultraviolet (UV)-visible light spectral absorbance of each ethanol wash supernatant. To check whether the pigments were fully removed from the algal tissue, we also measured the absorbance of the final fucoidan products. We compared the absorbance of our supernatants and fucoidan samples to the peak absorbance ranges of the major brown algal pigments: chlorophyll *a*, chlorophyll *c_2_*, and fucoxanthin, as measured in phytoplankton (Clementson & Wojtasiewicz, 2019). Brown algae also contain the pigment chlorophyll *c_1_*, but since chlorophyll *c_1_* has an absorbance spectrum that overlaps exactly with that of chlorophyll *c_2_* with relatively lower absorbance values (Jeffrey, 1969), the absorbance of chlorophyll *c_2_*obscures that of chlorophyll *c_1_*. In addition to pigments, brown algae contain phlorotannins, which are a type of polyphenol that absorb ultraviolet light. We also compared the total absorbance of the supernatants to the peak absorbance range of phlorotannins, as measured in *Sargassum linifolium* (Gheda et al., 2023).

The raw absorbance values of the ethanol wash supernatants and fucoidan samples cannot be directly compared, because the aliquots that were measured represent different proportions of the total sample material. To estimate the absorbance of the whole samples, we scaled the absorbance measurement to account for the entire volume of ethanol used for the wash steps or the entire mass of fucoidan yielded by the subsequent extraction steps (see Methods).

The absorbance spectra of the ethanol supernatants from the E100 and E70E100 wash schemes contain a significant peak at ∼315 nm (Figure 3A), indicating that phlorotannins were removed from the samples during the wash steps. The E70E100 supernatants had higher absorbance values surrounding this range than the E100 supernatants, indicating better removal of phlorotannins by the E70E100 wash scheme, consistent with the drop in polyphenols in the AIR samples seen previously (Figure 2G). Both supernatant spectra also contained peaks at wavelengths corresponding to the pigments chlorophyll *a* (380-450 and 640-680 nm), fucoxanthin (480-540 nm), and chlorophyll *c_2_* (570-640 nm) (Figure 3A). The absorbance of the E70E100 supernatants was higher in the 300-450 nm range compared to the absorbance of the E100 supernatants, indicating that the inclusion of 70% ethanol washes improved the removal of chlorophyll *a* (Figure 3A). The other peaks were of roughly equal absorbance values, indicating that chlorophyll *c_2_* and fucoxanthin removal was similar between the wash schemes. Overall, the absorbance data of the ethanol supernatants demonstrate that both wash schemes removed pigments and phlorotannins, but the E70E100 wash scheme continued to remove impurities better than the E100 wash scheme, consistent with the higher purity of the E70E100 AIRs.

**Figure 3.**
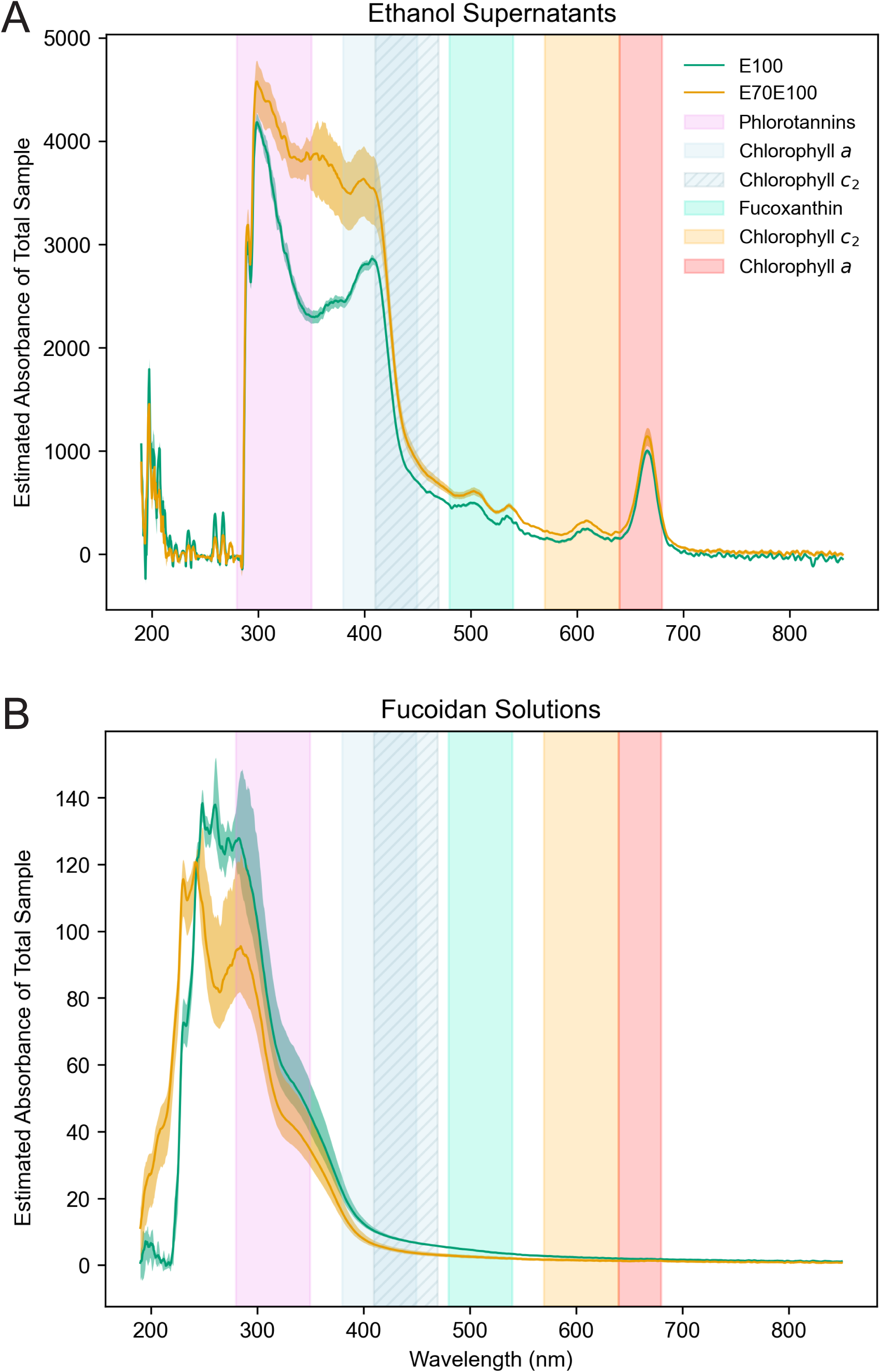
E70E100 wash scheme removes more chlorophyll a and phlorotannins. A) The total UV-visible light spectral absorbance of the ethanol supernatants from each of the wash steps that were performed to prepare the fucoidan samples E70E100 and E100. The sums of the absorbance values of the ten washes were calculated for each replicate independently (n=3). B) Absorbance of the fucoidan solutions prepared from samples E70E100 and E100. The peak absorbance ranges of phlorotannin (Gheda et al., 2023) and the three major pigments found in brown algae (chlorophyll *a*, chlorophyll *c_2_*, and fucoxanthin) (Clementson & Wojtasiewicz, 2019) are shaded. For A and B, Absorbance values of sample aliquots were multiplied by a scale factor to estimate the absorbance of the whole sample. The mean value of the replicates is plotted as a line, and the range from the minimum to maximum values for the replicates is shaded (n=3).

When the absorbance of the ethanol supernatants from washes 1-5 and 6-10 were viewed separately, it became apparent that 100% ethanol was primarily responsible for the removal of chlorophyll *a* (Supplemental Figure 1A and B). The absorbance of E100 washes 1-5 (100% ethanol) peaked at wavelengths corresponding to the absorbance of chlorophyll *a,* higher than the corresponding peaks for the E70E100 washes 1-5 (70% ethanol; Supplemental Figure 1A). The subsequent switch to 100% ethanol for E70E100 washes 6-10 resulted in noticeable peaks at the wavelengths corresponding to the red and blue absorbance of chlorophyll *a*, which was not observed in the E100 washes 6-10 (Supplemental Figure 1B). Although 100% ethanol appeared to be primarily responsible for the removal of chlorophyll *a*, the inclusion of 70% ethanol washes in the E70E100 wash scheme did result in higher overall absorbance at the wavelengths corresponding to chlorophyll *a*, indicating that combining 100% ethanol washes with 70% ethanol washes allows for more complete removal of chlorophyll *a*.

Next, we measured the UV-visible light spectral absorbance of the final fucoidan products to detect whether any pigments remained after the wash steps. In DNA extraction protocols, the A260/A230 ratio is often used a measure of purity, where DNA absorbs light at 260 nm, and any absorbance of light at 230 nm is commonly taken to indicate the presence of carbohydrates, polyphenols, lipids, and/or salts. Thus, we expected fucoidans to absorb light at 230 nm, because they are carbohydrates. We did not expect fucoidan molecules to absorb elsewhere in the UV-visible light spectrum, because the molecules do not have any light-absorbing structural features (such as aromatic rings). Indeed, the fucoidan samples absorbed light at around 230 nm (Figure 3B) while the ethanol supernatants did not (Figure 3A). A decrease in absorbance can be seen at 260 nm in the E70E100 fucoidan sample, but not in the E100 fucoidan sample, likely due to better removal of nucleic acids by the E70E100 wash scheme (Figure 3B).

Both E100 and E70E100 fucoidans have negligible absorbance at wavelengths of 380-800 nm, indicating that both wash schemes effectively removed all pigments (Figure 3B). In the regions corresponding to blue light absorbance by chlorophyll *a* and *c*_2_, the E70E100 sample had a slightly lower absorbance than the E100 sample, while the E70E100 ethanol supernatants have more absorbance (Figure 3A), providing more evidence that the E70E100 wash scheme removed more chlorophyll *a* and *c*_2_ than the E100 wash scheme. There was also high absorbance by the fucoidan samples in the region corresponding to phlorotannins (Figure 3B), indicating that not all phlorotannins were removed by the ethanol washes. The E100 fucoidan had higher absorbance in the phlorotannin region than the E70E100 fucoidan (Figure 3B), in accordance with the higher absorbance by the E70E100 ethanol supernatants in this range (Figure 3A). These results help confirm that the E70E100 wash scheme removed more phlorotannins, in agreement with the results of the biochemical polyphenol assay (Figure 2G). Overall, the UV-visible light absorbance data provide further evidence that the E70E100 wash scheme is superior for impurity removal.

### Additional solvents are unnecessary for impurity removal

Besides ethanol, common AIR extraction solvents include various combinations of acetone, methanol, and chloroform. Torode et al. followed their ethanol washes with acetone washes and then chloroform:methanol (3:2, v/v) washes (Torode et al., 2015). We wondered whether acetone or chloroform-methanol remove additional impurities that are not removed by ethanol washes. We started with our best performing wash scheme, E70E100, and continued with two additional wash schemes: 3 acetone washes (E70E100-A) or 3 chloroform:methanol (3:2 v/v) solution washes (E70E100-C; Figure 4A). We were particularly interested to see whether these other solvents would improve the removal of phlorotannins, since the E70E100 wash scheme did not fully remove phlorotannins (Figure 3). Neither additional solvent affected the AIR yield (Figure 4B) or the fucoidan yield (Figure 4C), indicating that they had little effect on fucoidan retention and removal of impurities. Indeed, the L-fucose and sulfate content was roughly equal in fucoidans from all three samples (Figure 4D and E), indicating that fucoidans were not lost during the additional solvent washes. The protein, polyphenol, and lipid content were also roughly equal in fucoidans from all three samples (Figure 4F-H), indicating that neither additional solvent provided additional impurity removal. Taken together, these data demonstrate that the addition of acetone or chloroform:methanol solution washes did not improve the purity of the fucoidan product.

**Figure 4.**
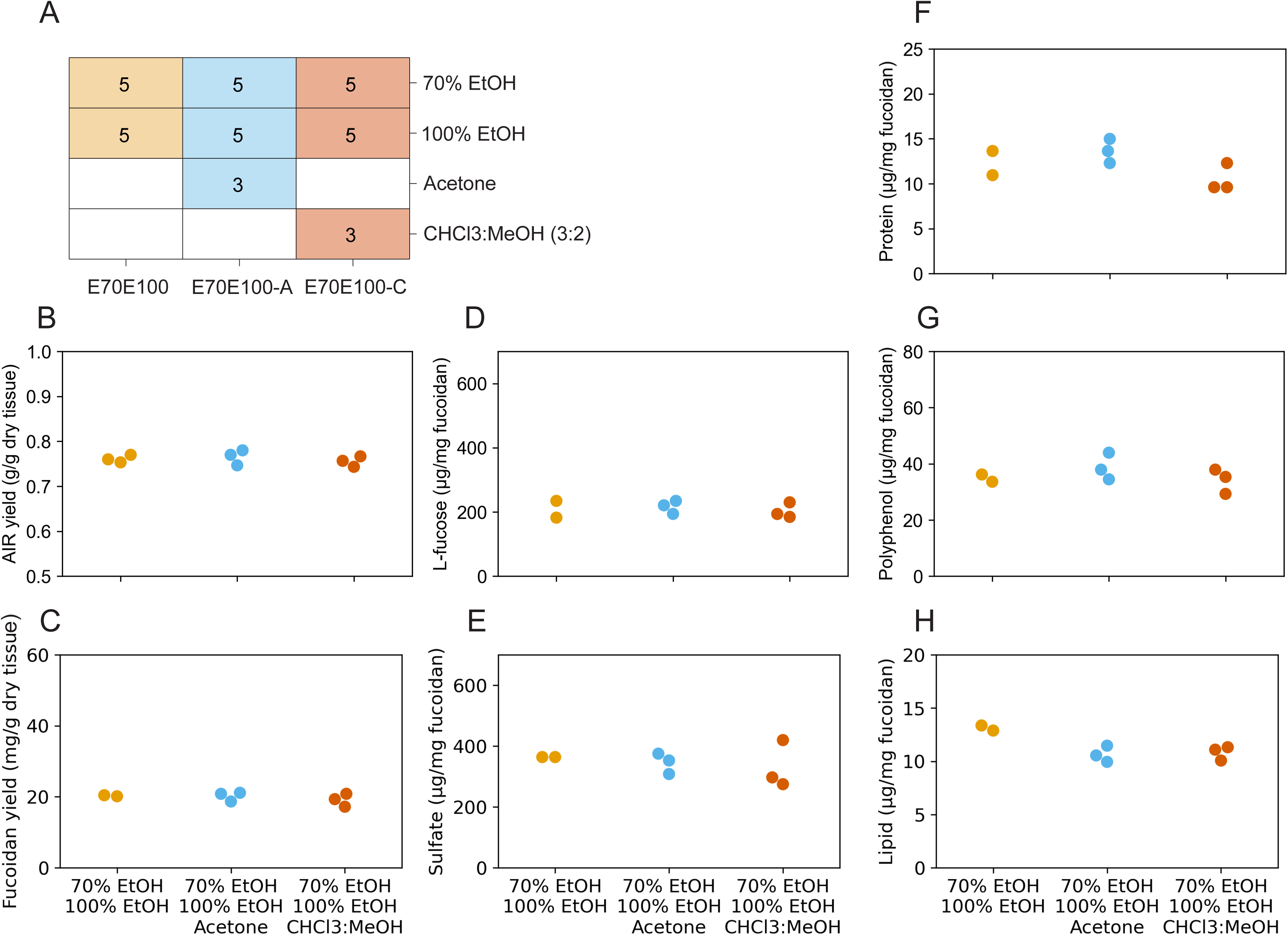
Additional solvents do not improve impurity removal from fucoidans. A) Schematic of additional solvent washes. The number inside a shaded box indicates the number of 5-minute washes that were conducted with the solvent indicated. B) AIR mass per gram of dry tissue for each wash scheme. C) Fucoidan mass per gram of dry tissue for each wash scheme. D) L-fucose content per mg of total fucoidan from each wash scheme as measured by an enzymatic L-fucose assay kit (MegaZyme). E) Sulfate content per mg of fucoidan from each wash scheme as measured by the Azure A assay. F) Protein contamination per mg of fucoidan as measured by the Bradford assay. G) Polyphenol contamination per mg of fucoidan as measured by the Folin-Ciolcateu assay. H) Lipid contamination per mg of fucoidan as measured by the sulfo-phospho-vanillin assay. The Mann-Whitney-Wilcoxon test two-sided was performed with p ≤ 0.05, and no statistically significant differences were found.

### Alginate is separated during fucoidan extraction with calcium chloride but not water

The next step in fucoidan extraction is to separate the fucoidans from other cell wall polysaccharides in the AIR, of which alginate is the most abundant. Water extractions are common, but CaCl_2_ solutions are often used instead (Dobrinčić et al., 2020), because calcium ions are expected to cross-link and insolubilize the alginates, separating them from the water-soluble fucoidans. However, alginate removal by extraction with CaCl_2_ solution has not been conclusively demonstrated. We hypothesized that fucoidans achieved through extraction in water would contain more alginate than fucoidans achieved through extraction in CaCl_2_ solution.

We performed fucoidan extractions from 2 g of E70E100 AIR in either water or 2% CaCl_2_. For each extraction solution, two volumes were tested (20 ml and 40 ml). All subsequent steps of the extraction protocol were identical between the samples. Following lyophilization, the mass of fucoidans was roughly equal in the samples extracted in 20 ml and 40 ml of CaCl_2_ solution (Table 1), indicating that the smaller volume was sufficient to extract most of the fucoidans from the AIR. Conversely, the samples extracted in 40 ml of water resulted in a fucoidan mass roughly three times higher than the samples extracted in 20 ml of either solvent (Table 1), indicating that water extracts components from the AIR that are not extracted by CaCl_2_ solution. We suspected that this increase in mass was due to co-extracted alginate rather than higher fucoidan yield.

**Table 1.** Fucoidan mass resulting from extraction using different solvent types and volumes. Identical AIR samples were extracted using the indicated solvent type and volume. Samples were centrifuged, dialyzed, and lyophilized. The mass of the final product was measured.

| Extraction solvent type | Extraction solvent volume (ml) | Fucoidan mass (mg) |
|---|---|---|
| H<sub>2</sub>O | 20 | 72.8 |
| H<sub>2</sub>O | 40 | 221.0 |
| 2% CaCl<sub>2</sub> solution | 20 | 68.1 |
| 2% CaCl<sub>2</sub> solution | 40 | 74.7 |

To detect the presence of alginate epitopes in the samples, we performed an enzyme-assisted immunosorbent assay (ELISA) with an alginate-directed brown algal monoclonal (BAM) antibody (Torode et al., 2016). We selected BAM10 in particular because it has demonstrated the highest sensitivity to multiple alginate epitopes (Torode et al., 2016). Alginate epitope signal was relatively higher in the positive control (alginic acid sodium salt from Sigma-Aldrich) and low in the negative controls (fucoidans and laminarin from Sigma-Aldrich; Table 2), demonstrating that the antibody was performing as expected.

**Table 2.**
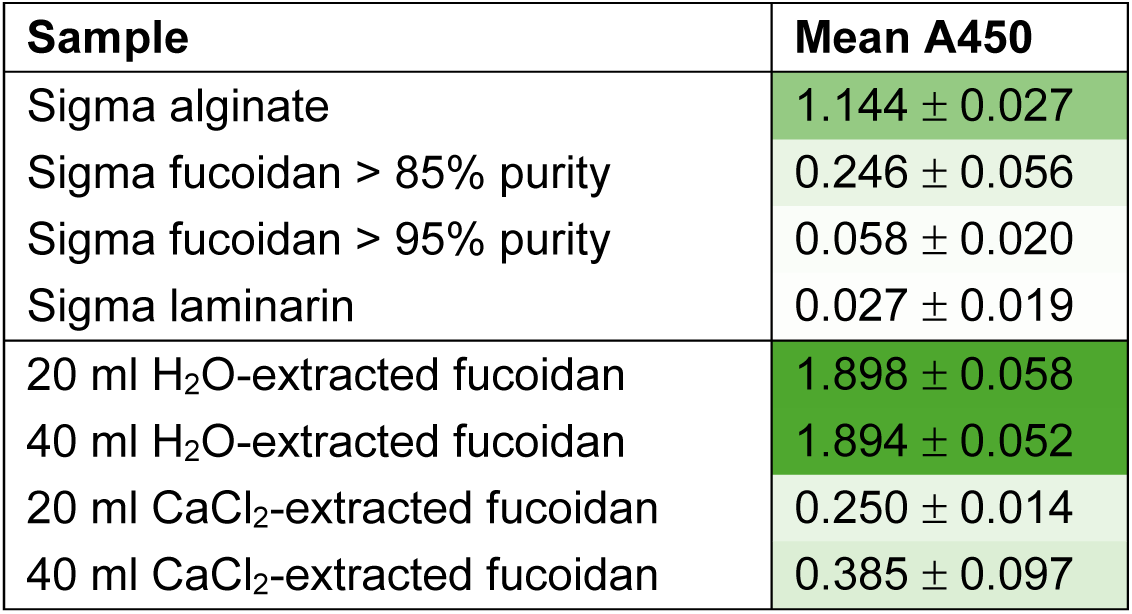
Detection of an alginate epitope in fucoidan extracts and commercial brown algal polysaccharides. An ELISA was performed with the alginate-directed brown algal monoclonal antibody 10 (BAM10; Torode et al., 2016) to detect the presence of an alginate epitope in the samples. The absorbance values are averages of biological replicates (n=3) with standard deviation. Each biological replicate value is the average of 3 technical replicates. The green color intensity scales from 0 to the highest value in the table.

Signal was higher in the fucoidan of >85% purity than the fucoidan of >95% purity (Table 2), suggesting that alginate accounts for some of the contamination in the less pure sample. Our fucoidan extracted in CaCl_2_ solution had slightly higher signal than the commercial fucoidan of >85% purity (Table 2), suggesting that the purity of our sample may be slightly lower.

Most importantly, the highest signal was detected in our fucoidans extracted in water (Table 2), confirming that water extracts alginates and indicating that the higher mass of the water-extracted samples was indeed due to higher alginate content rather than increased fucoidan yield. Thus, these data indicate that extraction in CaCl_2_ solution results in a purer fucoidan product with less co-extracted alginate. Since alginate makes up a considerable portion of the cell wall mass (Mabeau & Kloareg, 1987), the removal of alginate by CaCl_2_ solution is likely responsible for the considerable decrease in mass from AIR to fucoidan (Figure 2C and Figure 4C).

Another alginate-directed antibody, BAM8, was included in the ELISA alongside BAM10. BAM7-9 have indistinguishable epitope selectivity and detection levels (Torode et al., 2016) and we chose to use BAM8 as a representative. The results demonstrated the same trends (Supplemental Table 16).

### Fucoidans are lost through high-MWCO dialysis membranes

After CaCl_2_ extraction, the fucoidan solution must be dialyzed to remove residual CaCl_2_ that did not bind to alginates (Figure 1 step 6). Dialysis membranes are available in a range of molecular weight cut-offs (MWCO), but MWCO is rarely specified in the literature. We hypothesized that low-MW fucoidans would be lost during dialysis using a membrane with a higher MWCO. We took a CaCl_2_ extract supernatant and performed dialysis with membranes of four different MWCOs: 1.0, 3.5, 6-8, and 12.4 kDa. We then lyophilized each sample to obtain fucoidans. Solutions of the fucoidans were separated by agarose gel electrophoresis and visualized by staining with toluidine blue, which binds anionic polysaccharides, to test for differences in fucoidan retention by the different membranes.

We expected to see a decrease in dye signal at the bottom of the gel for samples dialyzed with higher MWCO membranes, as this would indicate that low-MW fucoidans were lost. Indeed, signal decreased in the samples dialyzed with the 12.4 kDa MWCO membrane; however, the decrease in signal was fairly uniform along the length of the lanes (Figure 5A). The intensity of the dye signal was quantified along the length of each lane, confirming that there was less signal in the 12.4 kDa-dialyzed samples across the length of the lane (Figure 5B). The 3.5 kDa-dialyzed samples bound the most dye across most of the length of the gel, indicating that this membrane retained the most fucoidans. The 6-8 kDa-dialyzed samples bound slightly less dye than the 3.5-dialyzed samples, except at the top of the gel where they bound the most dye of all the membranes. Surprisingly, the 1 kDa-dialyzed samples bound less dye than the 3.5 and 6-8 kDa-dialyzed samples. Together, these data indicate that fucoidans were lost during dialysis with higher MWCO (12.4 kDa) membranes.

**Figure 5.**
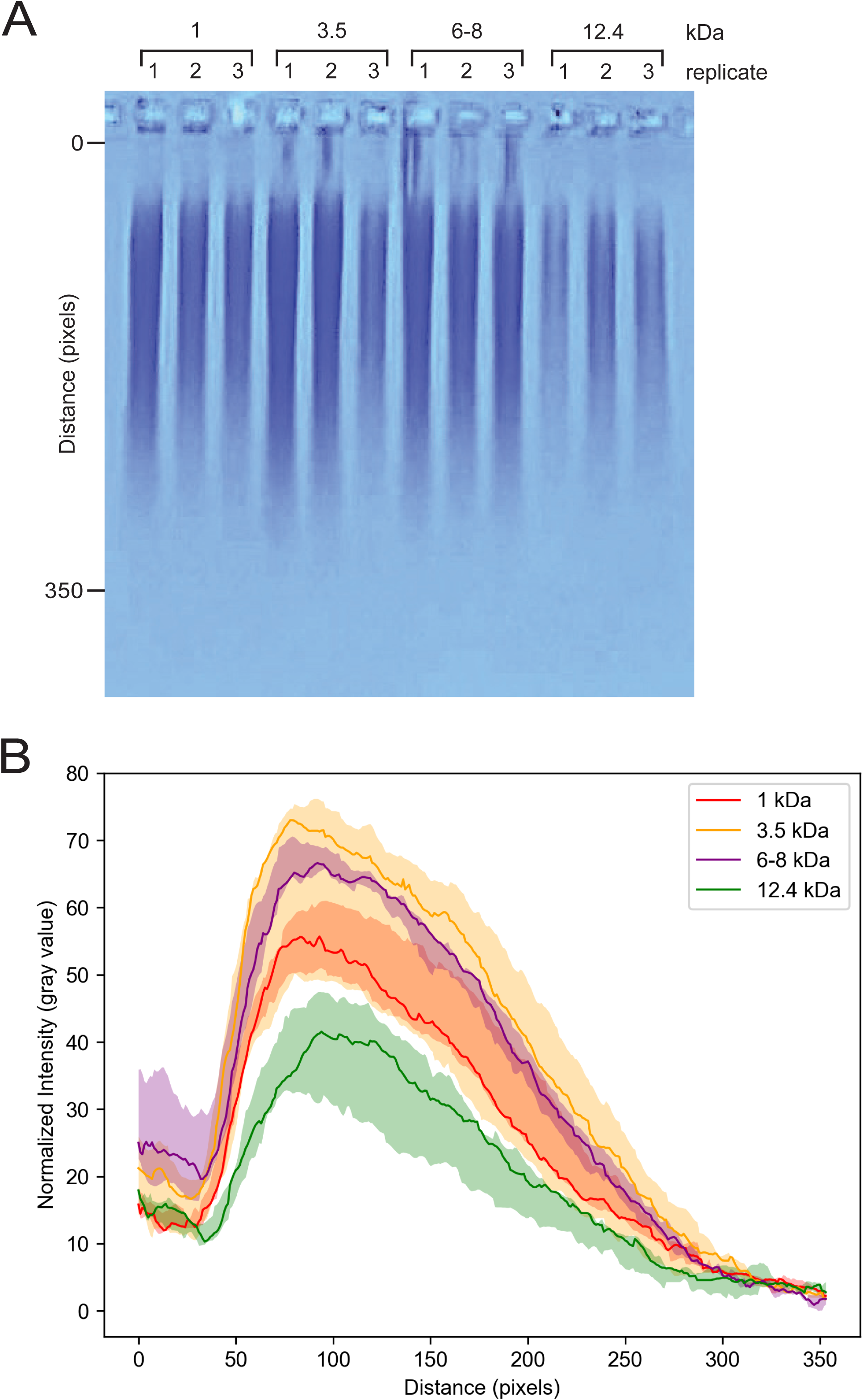
Low-MWCO dialysis membranes retain more fucoidans. A) A CaCl_2_ extract supernatant was dialyzed with tubing of the indicated MWCOs. Triplicate samples were separated by gel electrophoresis and stained with toluidine blue. B) The intensity of the dye signal along the lanes of the gel, starting just under the well, was quantified using Fiji (Schindelin et al., 2012). The region between the minimum and maximum value for each sample type is shaded, and the line indicates the median value (n=3).

## DISCUSSION

Fucoidans are extracted from brown algal tissue for both scientific investigations and industrial applications. Fucoidan extraction techniques are known to affect fucoidan composition and bioactivity, but extraction protocols are not standardized (Ale et al., 2011; Ko et al., 2026). Here, we identified multiple steps of a common extraction protocol as sources of variation between studies, and optimized each to maximize fucoidan yield and purity and overall protocol safety (Figure 1). The brown alga *S. compressa* was used in the experiments described here because it is abundant on the Southern California coast. However, we believe our findings would be applicable to many brown algal species due to similar overall cell wall architecture.

### Solvent selection for AIR extraction

First, we investigated impurity removal by different combinations of solvents during AIR extraction. We found that washing tissue with 70% ethanol and then 100% ethanol removed more proteins, polyphenols, and pigments than washing with an equal number of 100% ethanol washes (Figures 2 and 3). Thus, we recommend that AIR extractions be performed with subsequently increasing concentrations of ethanol. Our data indicate that the ethanol washes may leave behind some contaminating phlorotannins. It is possible that other less traditional solvents may more effectively remove phlorotannins, but we found that additional acetone or chloroform-methanol washes after the 70% and 100% ethanol washes did not remove additional impurities (Figure 4). In a similar investigation, Dobrinčić et al. found that washing tissue with both acetone and ethanol achieved better pigment removal than an equal number of washes with only 80% or 96% ethanol (Dobrinčić et al., 2021). It is possible that our E70E100-A wash scheme removed more pigments than our E70E100 wash scheme, but that was not measured here because acetone is not compatible with our NanoDrop One^C^. The addition of acetone washes may remove slightly more pigments, but 70% and 100% ethanol remove the majority of contaminants including most pigments.

### Solvent selection for fucoidan extraction

Second, we investigated the fucoidan extraction solution. Though acidic solutions and heat are often found to increase fucoidan yield (Hahn et al., 2012), acid extractions produce degraded fucoidan products (Ale et al., 2012) and high temperatures are expected to do the same. Recently, some less conventional fucoidan extraction technologies have emerged: ultrasound-, microwave-, and enzyme-assisted extractions, and pressurized liquid extractions. These are referred to as advanced extraction technologies, as they are more eco-friendly and produce higher yields of purer fucoidan products. However, all of them degrade the fucoidan product (Garcia-Vaquero et al., 2017). Since one of our aims was to extract fucoidan structures that are as close as possible to the native structures naturally occurring in the alga, we sacrificed a potentially higher fucoidan yield and extracted without acid, heat, or advanced technologies. This left two options: water or CaCl_2_ solution. Until a total alginate quantification method is developed, the detection of alginate epitopes using monoclonal antibodies is the best proxy assay for alginate quantification. We demonstrated that water extracts more mass than 2% CaCl_2_ solution (Table 1), and demonstrated that two alginate epitopes are significantly higher in the fucoidans extracted by water (Table 2, Supplemental Table 16), indicating that the extra mass in the water-extracted fucoidan is co-extracted alginate. Thus, we recommend extracting fucoidans with CaCl_2_ solution. Since there was little difference in the fucoidan mass extracted from 2 g of AIR by 20 ml and 40 ml of 2% CaCl_2_ solution, we recommend the lower volume (10 ml 2% CaCl_2_ solution per 1 g AIR) to save reagents.

For some applications, it may be useful to remember that the fucoidans extracted by aqueous solutions do not represent all of the fucoidans in the algal tissue; there are additional fucoidans that are more tightly attached to other cell wall components (KLOAREG & QUATRANO, 1988). Following extraction of water-soluble fucoidans from AIR in 2% CaCl_2_ solution, Torode et al. next solubilized alginates in alkaline solution (3% NaCO_3_). ELISA with fucoidan-directed antibodies showed that the NaCO_3_ extract gave higher signal than the CaCl_2_ extract, indicating that there are more alginate-bound fucoidans than water-soluble fucoidans (Torode et al., 2015). Since our goal was to obtain fucoidans free of co-extracted impurities, the data presented in this work only represent the fucoidan yield and composition of water-soluble fucoidans.

### MWCO selection for dialysis

Third, we investigated the effect of the MWCO of dialysis membranes on fucoidan retention. Toluidine blue binds to negatively charged high-MW polysaccharides, including alginate which is carboxylated and fucoidan which is sulfated (McCully, 1966). Since alginate was already shown to be removed from the fucoidan samples by the ELISA, we assume that the dye is only binding to fucoidans in the samples. We predicted that low-MW fucoidans would be lost during dialysis with higher MWCO membranes and predicted this would be visualized by decreased toludine blue signal in the lower end of a lane. Interestingly, we observed that increasing the membrane MWCO from 3.5 to 6-8 to 12.4 kDa resulted in decreased toluidine blue signal across the entire lane, not just the lower end (Figure 5).

We concluded that fucoidans were retained by the 3.5 kDa MWCO membranes, and increasing the MWCO resulted in loss of fucoidans; however, given the unexpected loss of total toluidine blue staining, we cannot say with certainty that it was only the low-MW fucoidans that were lost. It is possible that charge affects gel migration in addition to MW; perhaps smaller fucoidans that were lost were varied in their degree of sulfation, and therefore were spread throughout the lane of the gel rather than concentrated near the bottom. It is not clear why the 1 kDa MWCO membrane retained slightly less fucoidans than the 3.5 kDa MWCO membrane.

In studies investigating the MW range of fucoidans using size-exclusion chromatography, fucoidans are often obtained through extractions involving dialysis with high or unspecified MWCO membranes (ex: (Gómez-Ordóñez et al., 2012; Neupane et al., 2020)), potentially obscuring the lowest MW fucoidans from the reported MW range. In other words, the fucoidan MW ranges reported in such studies may have an artificially high lower bound. Our data emphasize the importance of selecting and reporting the membrane MWCO used for dialysis in studies aimed at understanding fucoidan MW ranges or optimizing fucoidan extraction yield.

### Other cell wall polysaccharides

The fucoidan extraction protocol used in this work does not have an obvious step where laminarin and cellulose are removed. Laminarins are 1-7 kDa (Spicer et al., 2017) and might be removed during dialysis if the MWCO is set for higher than laminarin MW but lower than fucoidan MW. However, it would be difficult to determine this threshold, because dialysis is needed to obtain the polysaccharides in order to perform MW analyses. Laminarins are made up of β-1,3-linked glucose monomers. Monomer and linkage analyses of our fucoidan samples reveal that there is glucose in our fucoidan samples, but not linked at the third carbon (data not shown), indicating that the glucose in our sample is part of a heterogenous fucoidan molecule rather than laminarin. We suspect that laminarin was removed at some step of the fucoidan extraction protocol, but it is not clear at which step. Brown algal cellulose has an average MW of 269 kDa, which would certainly be retained with fucoidans during dialysis if not removed earlier in the extraction process. Cellulose is made up of β-1,4-linked glucose monomers, which may also be present in fucoidans, making it difficult to determine whether a fucoidan sample contains cellulose.

## CONCLUSION

Our analyses examined multiple steps of a typical fucoidan extraction method. Our data indicate that a combination of 70% and 100% ethanol washes alone removes most impurities during the preparation of the AIR; thus, the most effective solvent option the safest and most eco-friendly as well. We also demonstrate that extracting fucoidans with CaCl_2_ solution results in far less alginate contamination than extracting with water, and recommend an AIR:solvent ratio for fucoidan extraction. Finally, we showed that the MWCO of dialysis membranes should be chosen carefully, as fucoidans are lost during dialysis with higher MWCO membranes.

Altogether, we hope that these findings help the field move toward a more standardized fucoidan extraction approach, to enable better comparisons of data from different studies. Extracting higher yields of purer fucoidans will help studies of fucoidan bioactivities in medicine and agriculture, and obtaining fucoidan structures which are more similar to the structures occurring *in vivo* will help studies of the function of fucoidan in brown algae and microbial degradation of fucoidans.

## DATA AVAILABILITY

All data can be found in the Supplemental materials.

## AUTHOR CONTRIBUTIONS

Conceptualization - JCK, SAB

Investigation - JCK

Formal Analysis – JCK

Visualization - JCK

Writing – original draft - JCK

Writing – review & editing - SAB, LKD

Funding Acquisition - SAB

Project Administration - JCK, LKD, SAB

Resources - LKD

Supervision - SAB, LKD

## ACKNOWLEDGEMENTS

Thank you to Dr. Stuart Conway, Dr. Glen Brodie, and Daniela Prochazkova for access to their lyophilizer. Thank you to Dr. Robert Gunsalus for access to his 96-well plate reader. Thank you also to Hugh Knopp for help with C-PAGE.

## LIST OF ABBREVIATIONS

AIR: alcohol-insoluble residue
MWCO: molecular weight cut-off
BAM: brown algal monoclonal antibody
UV: ultraviolet
ELISA: enzyme-linked immunosorbent assay
C-PAGÈ: carbohydrate polyacrylamide gel electrophoresis

---

<u>Supplemental Figure 1. 100% ethanol is primarily responsible for the removal of chlorophyll *a*.</u>

A) Total absorbance of the ethanol supernatants from washes 1-5 for each wash scheme. The sums of the absorbance values for the five washes were calculated for each replicate independently (n=3). Absorbance values were normalized to the highest value observed between the two samples, which was set to 1.0. The mean value is plotted as a line, and the range from the minimum to maximum values for the three replicates is shaded. B) Total absorbance of the ethanol supernatants from washes 6-10 of the algal tissue samples. The sums of the absorbance values for the ten washes were calculated for each replicate independently (n=3). Absorbance values were normalized to the highest value observed in A, which was set to 1.0. The mean value is plotted as a line, and the range from the minimum to maximum values for the three replicates is shaded. C) Total absorbance of the ethanol supernatants from all ten E100 washes, and absorbance of a 15 mg/ml solution of the E100 fucoidan sample. D) Total absorbance of the ethanol supernatants from all E70E100 washes, and absorbance of a 15 mg/ml solution of the E70E100 fucoidan sample. Across C and D, wash and fucoidan absorbance values were normalized independently to the highest value observed in each respective dataset, which was set to 1.0. The mean value is plotted as a line, and the range from the minimum to maximum values for the three replicates is shaded.

<u>Supplemental Figure 2. Manual grinding stages of dried S. *compressa* tissue.</u> Tissue appearance after the first, second, third, and fourth round of grinding in liquid nitrogen, from left to right.

<u>Supplemental Figure 3. A monosulfated L-fucose monomer is 33% sulfate by mass.</u> A) A Fischer projection of L-fucose with sulfation at the 3rd carbon. Carbons are labeled by red numbers. The entire monosulfated L-fucose molecule is shaded in blue, and the sulfate group is shaded in orange. B) The individual and total masses of the elements in the sulfate group alone (orange) and the monosulfated L-fucose molecule (blue). C) The mass of the sulfate group makes up 33% of the mass of the monosulfated L-fucose molecule.

